# Allele-Resolved Hybrid Embryos Reveal the Fates of Regulatory Divergence

**DOI:** 10.64898/2026.09.11.750430

**Authors:** Jiali Lei, Laurence A. Lemaire, Hidehiko Hashimoto, Liam Rodman, Michael Levine, Chen Cao

**Affiliations:** Department of Bioengineering, University of Texas at Dallas, Richardson, TX, USA; Department of Biology, Saint Louis University, St. Louis, MO, USA; Lewis-Sigler Institute for Integrative Genomics, Princeton University, Princeton, NJ, USA; Department of Molecular Biology, Princeton University, Princeton, NJ, USA

## Abstract

Developmental programs can remain conserved despite extensive regulatory divergence, but how evolved regulatory differences are transmitted through embryonic lineages remains unclear. Here we generate a time resolved, allele resolved single-cell atlas of hybrid embryogenesis between *Ciona intestinalis* and *Ciona savignyi*, enabling regulatory differences accumulated between species to be followed across defined developmental lineages. We find that allelic differences are maintained or remodeled in lineage specific ways, with their outcomes associated with regulatory origin and allele specific chromatin accessibility. Across multiple tissues, allelic divergence increases along gene regulatory network (GRN) hierarchy from upstream regulators toward downstream regulators and effector genes. In the cardiopharyngeal lineage, *Foxf* illustrates how allelic dominance provides partial compensation for highly divergent regulatory sequences and thereby contributes to developmental system drift. Together, our results reveal that regulatory divergence is dynamically sorted during development according to lineage context and GRN hierarchy. This lineage resolved framework provides a developmental basis for understanding how extensive regulatory evolution can accumulate while conserved embryonic programs are maintained.

## Introduction

The evolution of animal form is largely driven by changes in the spatial and temporal regulation of gene expression. A central challenge in evolutionary developmental biology is to explain how extensive genotypic diversification can occur while preserving the robustness of embryonic body plans. Although cis-regulatory elements (CREs)^1,2^ are widely regarded as major drivers of evolutionary change because they enable tissue-specific modulation of gene expression, cross-species analyses have revealed that developmental programs can remain highly conserved despite extensive divergence in regulatory sequence^3,4^. This apparent paradox, commonly referred to as developmental system drift^5^, suggests that developmental stability does not require strict conservation of individual regulatory elements, but can emerge from higher-order organization of developmental regulation. One prevailing model proposes that gene regulatory networks (GRNs) are organized into semi-independent regulatory modules^5-7^, allowing evolutionary changes in individual modules to accumulate without disrupting core developmental processes. Consistent with this view, comparative studies have revealed both conserved developmental subcircuits and flexible regulatory components across animal systems^8,9^.

However, how such modular regulatory evolution is implemented within an intact developing embryo, and whether developmental context influences the outcome of regulatory divergence, remain poorly understood. Developmental system drift implies that GRNs can preserve stable outputs despite regulatory divergence, but this stability alone does not explain how embryos organize and accommodate such divergence. Recent single-cell hybrid and allele-specific expression studies have shown that cis- and trans- regulatory effects and allelic imbalance can vary across selected cell types and restricted differentiation trajectories^10-12^. Yet how allelic regulatory divergence is coordinated across branching embryonic lineages is not known, particularly as multiple natural lineages diversity and pass through distinct regulatory states. To address this question, we used hybrid embryos generated between the ascidians *Ciona intestinalis* and *Ciona savignyi*. Despite their highly similar larval body plans, these species are separated by approximately 180 million years of evolution^13^ and show extensive genomic divergence^14^. Both species also harbor unusually high levels of genetic variation within natural populations, even among marine invertebrates with large effective population sizes such as sea urchins^15,16^. More strikingly, orthologous sequences between *C. intestinalis* and *C. savignyi* differ by approximately 15%^17^, a level of divergence comparable to that between human and rodent orthologues. This extensive sequence divergence provides a dense landscape of species specific variants, enabling the two parental alleles to be distinguished with high confidence in hybrid embryos. Nevertheless, viable hybrids progress through normal embryogenesis. In these embryos, homologous alleles carrying divergent regulatory architectures experience the same cellular environment, allowing regulatory differences to be read out as allele-specific activity within matched developmental contexts. This combination of extensive sequence divergence and developmental compatibility therefore provides an unusual opportunity to follow evolutionarily diverged regulatory programs across an intact embryonic time course.

Leveraging this system, we generate a time-resolved, allele-resolved single-cell atlas of *Ciona* hybrid embryogenesis, integrating transcriptomic and chromatin accessibility measurements across embryonic lineages and stages. By following the activity of the two species derived alleles through development, we find that allelic regulatory divergence is not transmitted uniformly through embryogenesis. Instead, the outcomes of allelic differences vary among lineages and are associated with their regulatory origin and allele specific chromatin accessibility. Across multiple tissues, divergence also increases along GRN hierarchy from upstream regulators toward downstream regulators and target genes. Together, these findings suggest that development acts as an interpreter of regulatory evolution, determining whether divergent regulatory inputs are retained as lineage specific memory or remodeled during development. This relationship provides a developmental framework for understanding how regulatory divergence can accumulate while conserved embryonic programs are maintained.

## Results

### Hybrid embryos reveal heterogeneous allelic regulation across lineages

To define the developmental landscape of allelic regulation, we generated reciprocal interspecific hybrids between *Ciona intestinalis* and *Ciona savignyi* (Fig. 1a and Extended Data Fig. 1) and profiled hybrid embryos by allele-resolved single-cell transcriptomics across 10 developmental stages, from initial gastrula to larva (Fig. 1b). Morphological comparison showed that embryos generated with *C. intestinalis* eggs and *C. savignyi* sperm developed comparably to the corresponding intraspecific controls and reached larval stages at high frequency (∼89%), whereas the reciprocal cross using *C. savignyi* eggs and *C. intestinalis* sperm rarely produced larvae (∼7%) and failed predominantly after gastrulation (Extended Data Fig. 2 and Extended Data Table 1). We therefore used the viable *C. intestinalis* egg × *C. savignyi* sperm cross for the primary allele-resolved developmental atlas, while reciprocal-cross embryos were used to distinguish species-of-origin from parent-of-origin associated allelic biases (Methods and Extended Data Table 2). The hybrid scRNA-seq atlas recovered 70,301 cells and captured all major embryonic lineages, including central and peripheral nervous system, epidermis, endoderm, mesenchyme, muscle, heart and notochord (Fig. 1b and Extended Data Table 3). Notably, the atlas also recovered rare cell populations, such as germ cells, indicating sufficient sensitivity to resolve low-abundance embryonic states. Across these cells, we detected expressions from more than 10,000 orthologous genes (Methods, Extended Data Table 4). Cell-type identities, lineage composition and the relative abundance of major cell populations in viable hybrids closely resembled those observed in parental and published *C. intestinalis* embryos^18,19^, supporting the conclusion that overall developmental patterning is largely preserved in the hybrid background (Fig. 1b and Extended Data Fig. 3).

**Figure 1.**
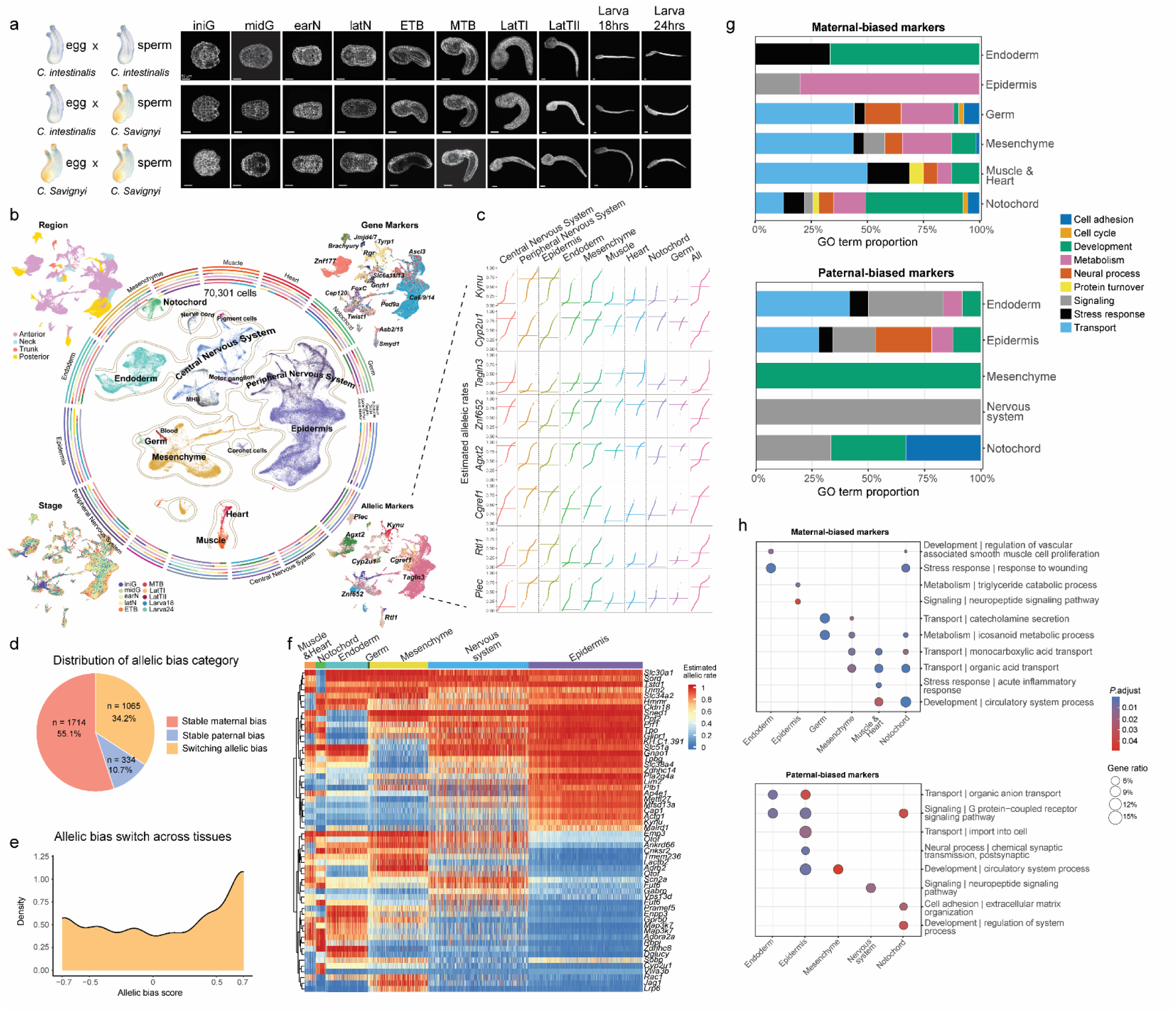
Allelic regulation is heterogeneous across cell types in the developing hybrid embryo. **a,** Schematic of the *C. intestinalis* and *C. savignyi* intraspecific control crosses and the *C. intestinalis* eggs × *C. savignyi* sperm interspecific hybrid cross used in this study (left panel). Representative embryos from the intraspecific (*C. intestinalis* egg × *C. intestinalis* sperm and *C. savignyi* egg × *C. savignyi* sperm) and hybrid (*C. intestinalis* egg × *C. savignyi* sperm) crosses across 10 developmental stages, from initial gastrula to larva (right panel). hrs, hours. iniG, initial gastrula; midG, mid-gastrula; earN, early neurula; latN, late neurula; ETB, early tailbud; MTB, mid-tailbud; LatTII, late tailbud II. Scale bars: 50 µm. **b,** Single-cell transcriptomic atlas of *C. intestinalis* egg × *C. savignyi* sperm hybrid embryos across ten developmental stages (n = 70,301 cells). Cells are annotated by major tissue and cell-type identities. Peripheral UMAPs show cells colored by anatomical region (upper left), developmental stage (lower left), expression of representative cell-type marker genes (upper right), and representative allelic marker patterns (lower right). Concentric annotation tracks indicate tissue, sub-tissue, developmental stage, anatomical region, gene-marker identity, and allelic-marker identity. n = 2 biological replicates at MTB stage; n = 3 biological replicates at LatTII stage. **c,** Estimated allelic-rate profiles of representative allelic marker genes across major tissues and cell types. **d,** Classification of genes according to the stability of allelic bias across cell types, including stable maternal bias, stable paternal bias, and switching allelic bias. Numbers and percentages indicate the genes assigned to each category. **e,** Distribution of allelic-bias scores across tissues for genes classified as exhibiting switching allelic bias. **f,** Heatmap of estimated allelic rates for the top 10 maternal- and paternal-biased marker genes across tissues. Rows represent allelic marker genes (n = 56 genes) and columns represent tissue identities; color denotes the estimated allelic rate, with 1 indicating exclusive *C. intestinalis* allele expression, 0 indicating exclusive *C. savignyi* allele expression, and 0.5 indicating balanced expression between the two alleles. Values above 0.5 indicate increasing bias toward the *C. intestinalis* allele, whereas values below 0.5 indicate increasing bias toward the *C. savignyi* allele. **g,** Functional composition of significantly enriched GO terms (Benjamini-Hochberg-adjusted *P* < 0.05) associated with maternal- and paternal-biased marker genes across tissues. GO terms are grouped into broad functional categories, and bar lengths indicate the proportional contribution of each category among enriched terms. Tissues without significantly enriched GO terms are not shown. **h,** Representative enriched GO terms associated with maternal- and paternal-biased marker genes across tissues. For each tissue, representative terms were selected from the two most abundant broad functional categories shown in **g**. Dot size indicates gene ratio, and color denotes the Benjamini-Hochberg-adjusted *P* value.

We next assigned allele-informative reads to the *C. intestinalis* or *C. savignyi* allele and quantified allelic imbalance within stage- and cell-type-specific contexts^20^ (Methods). Allelic rate ranges from 0 to 1, with higher values indicating greater *C. intestinalis* allele contribution, lower values indicating greater *C. savignyi* allele contribution, and 0.5 indicating balanced allelic expression. Canonical lineage markers identified in intraspecific *Ciona* embryos were recovered in the corresponding hybrid cell populations, supporting consistent annotation of major tissues and cell states^18^ (Fig. 1b and Extended Data Table 5). Biological replicates showed consistent major cell-type composition and allelic-bias patterns, supporting the reproducibility of the observed tissue-level allelic differences (Extended Data Fig. 4). Within these annotated contexts, we identified allele-biased marker genes for individual tissues and cell states (Fig. 1c and Extended Data Table 6). For instance, *Cyp2u1* appeared close to balanced when averaged across the embryo, but showed strong paternal bias in epidermis and peripheral nervous system. Conversely, *Cgref1* and *Agxt2* displayed pronounced tissue dependent variation in allelic bias, but shifted toward the opposite parental direction in epidermal derivatives. Additional genes, including *Kynu*, *Tagln3*, *Znf652*, *Rtl1* and *Plec*, showed similarly tissue-restricted allelic patterns. These examples indicate that allelic imbalance is shaped by tissue identity and cell state, and can be obscured by bulk averaging (Fig. 1c). To determine whether allelic bias was stable across cell types or changed in a context-dependent manner, we classified genes by the consistency of their bias direction (Methods). This analysis identified 1,714 stable maternal-biased genes, 334 stable paternal-biased genes and 1,065 switching genes (Fig. 1d). A signed consistency score further captured the extent of directional stability (Fig. 1e, Methods), indicating that recurrent maternal bias is the predominant stable allelic pattern across cell types. Heatmap analysis further revealed that allelic imbalance was organized into cell type associated modules rather than a uniform embryo-wide pattern, separating broad parental-bias programs from tissue restricted and cell state-specific allelic programs (Fig. 1f). Gene Ontology (GO) enrichment analysis showed that maternal- and paternal-biased markers were associated with distinct tissue-dependent functional programs (Fig. 1g,h and Extended Data Table 7). Maternal-biased markers were enriched for stress-response, metabolic and transport terms, whereas paternal-biased markers were enriched for signaling and extracellular matrix associated programs. These enrichments varied across tissues, indicating that allelic regulatory divergence is organized into lineage-specific functional modules rather than distributed uniformly across the embryo. Together, these data show that viable *Ciona* hybrid embryos preserve overall morphology and cell-type composition while exposing extensive, tissue-compartmentalized allelic divergence at single-cell resolution.

### Developmental lineage progression defines allelic memory and remodeling

We next asked how parental allelic activity is organized along the embryonic lineage tree. To relate allelic imbalance to developmental history, we projected allele-resolved expression states onto a time-resolved *Ciona* lineage framework^18^ (Methods), in which each node represents a defined cell population at a given developmental stage, and each edge represents developmental continuity between connected parent and child states (Fig. 2a and Extended Data Fig. 5). This representation allowed us to distinguish two levels of allelic organization: the allelic composition of each node, and the fate of allelic states as cells progress from one node to the next. After excluding maternally loaded and lowly expressed genes (Methods), we quantified maternal- and paternal-allele bias across the lineage tree. The allelic landscape changed progressively over developmental time, with the proportion of paternal biased genes increasing from early to later stages, while balanced and maternal biased fractions varied among lineages (Extended Data Fig. 6). The increase in paternal bias is consistent with progressive zygotic deployment of the paternal *C. savignyi* genome as embryonic transcription and lineage-specific regulatory programs become established^17^. Related lineages within the same broad tissue also showed distinct allelic compositions (Extended Data Fig. 7), further indicating that allelic imbalance is shaped not only by developmental time, but also by lineage identity and regulatory state.

**Figure 2.**
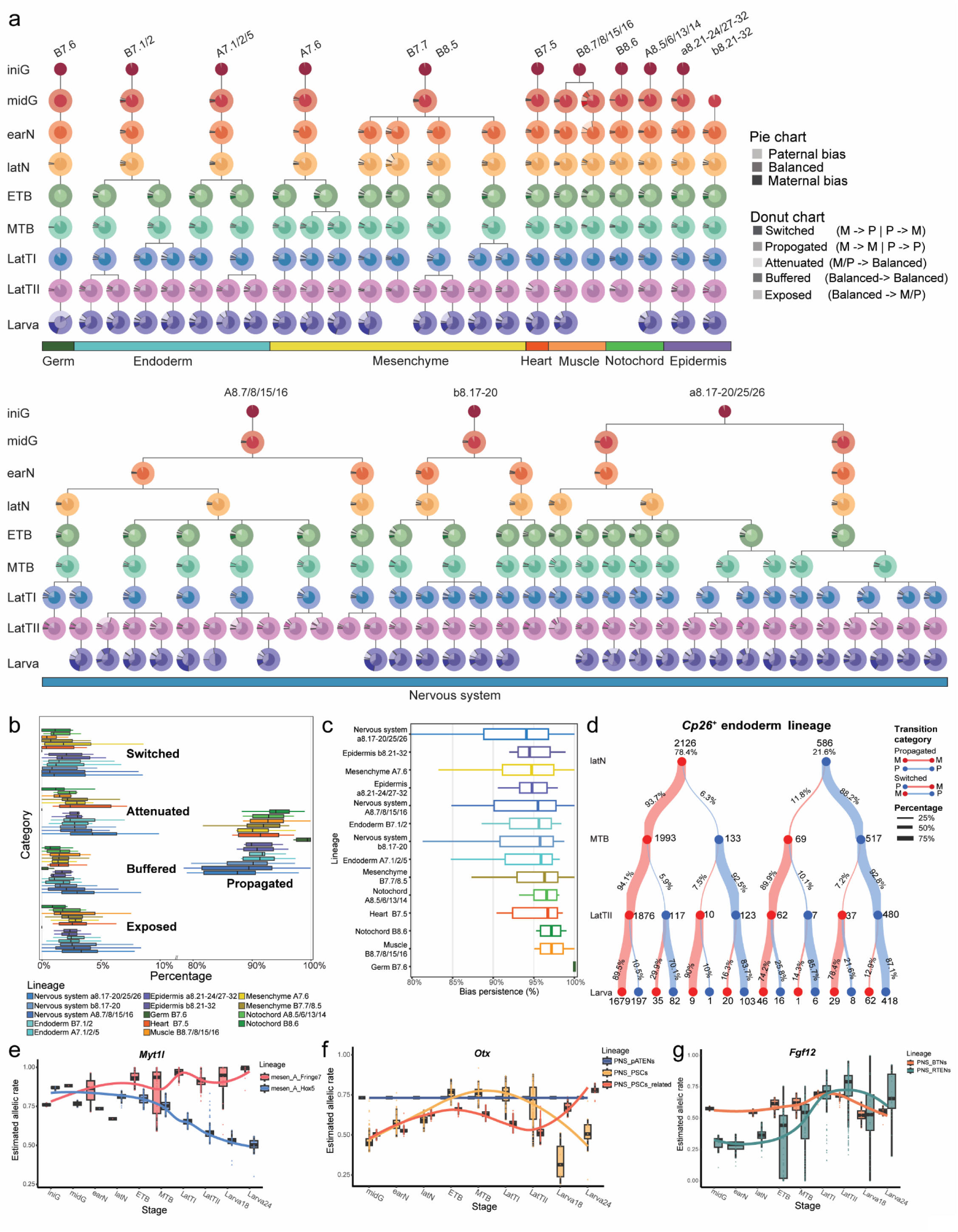
Lineage progression reveals distinct modes of allelic memory and remodeling. **a,** Lineage-resolved trajectories of allelic bias across embryonic development, from initial gastrula to larval stages (n = 64,788 cells). Pie charts show the proportions of genes classified as maternally biased, balanced, or paternally biased at each developmental state. Donut charts summarize allelic-state transitions between consecutive states within each lineage, classified as propagated (maternal to maternal or paternal to paternal), switched (maternal to paternal or paternal to maternal), attenuated (maternal/paternal to balanced), buffered (balanced to balanced), or exposed (balanced to maternal/paternal). Developmentally related cell states are connected according to the reconstructed lineage hierarchy, and node colors denote developmental stage. Major tissue identities are indicated below the trajectories. M, maternal bias; P, paternal bias. Missing nodes represent cell states containing fewer than six cells and were excluded from the analysis. **b,** Proportions of genes assigned to each allelic-state transition category across individual developmental lineages. Colors denote lineages. **c,** Distribution of allelic-bias persistence across developmental lineages. Bias persistence was defined, among transitions originating from an allelically biased state, as the proportion of propagated transitions relative to propagated, switched, and attenuated transitions. Boxplots summarize persistence values across developmental transitions within each lineage. **d,** Example of allelic-bias propagation and remodeling along the *Cp26*^+^ endoderm lineage from late neurula to larva. Red and blue nodes denote maternally and paternally biased gene states, respectively, and numbers indicate gene counts at each state. Connecting ribbons trace the fate of biased genes between consecutive developmental stages; ribbon width reflects the fraction of genes following each transition, with percentages shown along the ribbons. **e,f,g,** Dynamic changes in estimated allelic rates of *Myt1l*, *Otx* and *Fgf12* across sampled developmental stages in different cell types.

To determine whether node-level differences reflected remodeling of allelic states during lineage progression, we classified each parent-to-child edge by the direction of allelic state change. This analysis distinguished maintained balance, retained parental bias, newly acquired bias, loss of bias and reversal of parental direction, corresponding to buffered, propagated, exposed, attenuated and switched transitions, respectively (Fig. 2a and Extended Data Table. 8). By converting lineage relationships into directional allelic transitions, this edge-based framework allowed us to distinguish allelic memory from allelic remodeling during development. Quantification of these transition classes across lineages showed that propagation was the predominant fate of pre-existing allelic bias, whereas attenuation, switching and exposure occurred less frequently and varied across lineages (Fig. 2b and Extended Data Fig. 8). Thus, although allelic bias was generally retained during lineage progression, a substantial subset of genes underwent lineage-dependent remodeling. The same gene could maintain bias, acquire bias, lose bias or reverse parental direction depending on developmental context (Extended Data Fig. 9), indicating that allelic behavior is shaped by the lineage state in which a gene is deployed. Bias persistence analysis further revealed substantial lineage differences in allelic memory (Methods, Fig. 2c). Germ cells showed the strongest apparent bias persistence, potentially reflecting the prolonged influence of maternally inherited germ-plasm components and delayed remodeling of early germline transcriptional programs^21,22^, whereas several somatic lineages showed greater allelic turnover. For example, the sensory-vesicle lineage showed pronounced fluctuations in bias persistence across successive transitions, suggesting that changing neural regulatory environments repeatedly expose or attenuate allelic differences during differentiation. These differences indicate that lineage progression acts as a regulatory filter, with the degree of allelic memory or remodeling depending on developmental contexts.

To visualize allelic remodeling within a defined developmental trajectory, we examined the endoderm lineage using Sankey analysis across successive stages (Fig. 2d). Although many genes retained the same parental direction across adjacent stages, 23% underwent a change in allelic state during lineage progression, including transitions between balanced and biased states and switches in parental direction. This shows that even within a coherent lineage trajectory, allelic divergence is not simply propagated forward, but is selectively stabilized, attenuated or reconfigured as the lineage matures. Representative transcription-factor trajectories further illustrate this context dependence. Mesenchyme-associated regulators such as *Myt1l* and *Meis1* showed branch-specific persistence or attenuation, whereas neural regulators including *Smad1*, *Fgf12* and *Otx* followed distinct temporal trajectories across related nervous-system lineages (Fig. 2e-g, Extended Data Fig. 10). Together, these analyses show that developmental lineages do not passively transmit a fixed allelic program. Instead, lineage progression reshapes the developmental expression of regulatory divergence. Developmental trajectories may therefore determine where and when regulatory differences accumulated between species emerge during embryogenesis.

### Regulatory architecture informs allelic fate during lineage progression

The lineage-transition analysis defined distinct allelic fates. To determine whether these fates were linked to regulatory architecture, we classified the sources of allelic imbalance in the hybrid embryo. Allelic bias in a hybrid embryo can arise from species-specific, allele-intrinsic cis-regulatory divergence or parent-of-origin associated effects, whereas trans-regulatory divergence can be inferred by comparing expression differences between the two intraspecific controls with allelic imbalance in the hybrid^11,17,23,24^. Here we treat “parent-of-origin associated” as an operational classification rather than evidence of genomic imprinting, because reciprocal cross asymmetries may also arise from maternal cytoplasmic factors or from interactions between species divergent cis sequences and the maternal trans regulatory environment^17,24^. We therefore integrated reciprocal crosses, intraspecific controls and allele-resolved chromatin accessibility to assign regulatory sources and test how they were distributed across lineage-transition fates (Methods). We first used allele-resolved single-cell transcriptomes from reciprocal crosses at the mid-gastrula stage to distinguish species-specific from parent-of-origin-associated biases (Fig. 3a). Maternally loaded transcripts were excluded, so these classifications reflect zygotically expressed allelic patterns rather than direct carryover of deposited maternal RNA. Most classified biases were species-specific, accounting for the largest fractions of biased genes. Parent-of-origin associated biases formed a smaller but substantial layer, dominated by maternal-origin associated patterns, whereas paternal-origin associated biases were rare. Consistent with this classification, variance partitioning analysis at mid-gastrula showed that species identity explained the largest fraction of allelic-rate variation, with parent of origin contributing a substantial secondary component (Fig. 3b).

**Figure 3.**
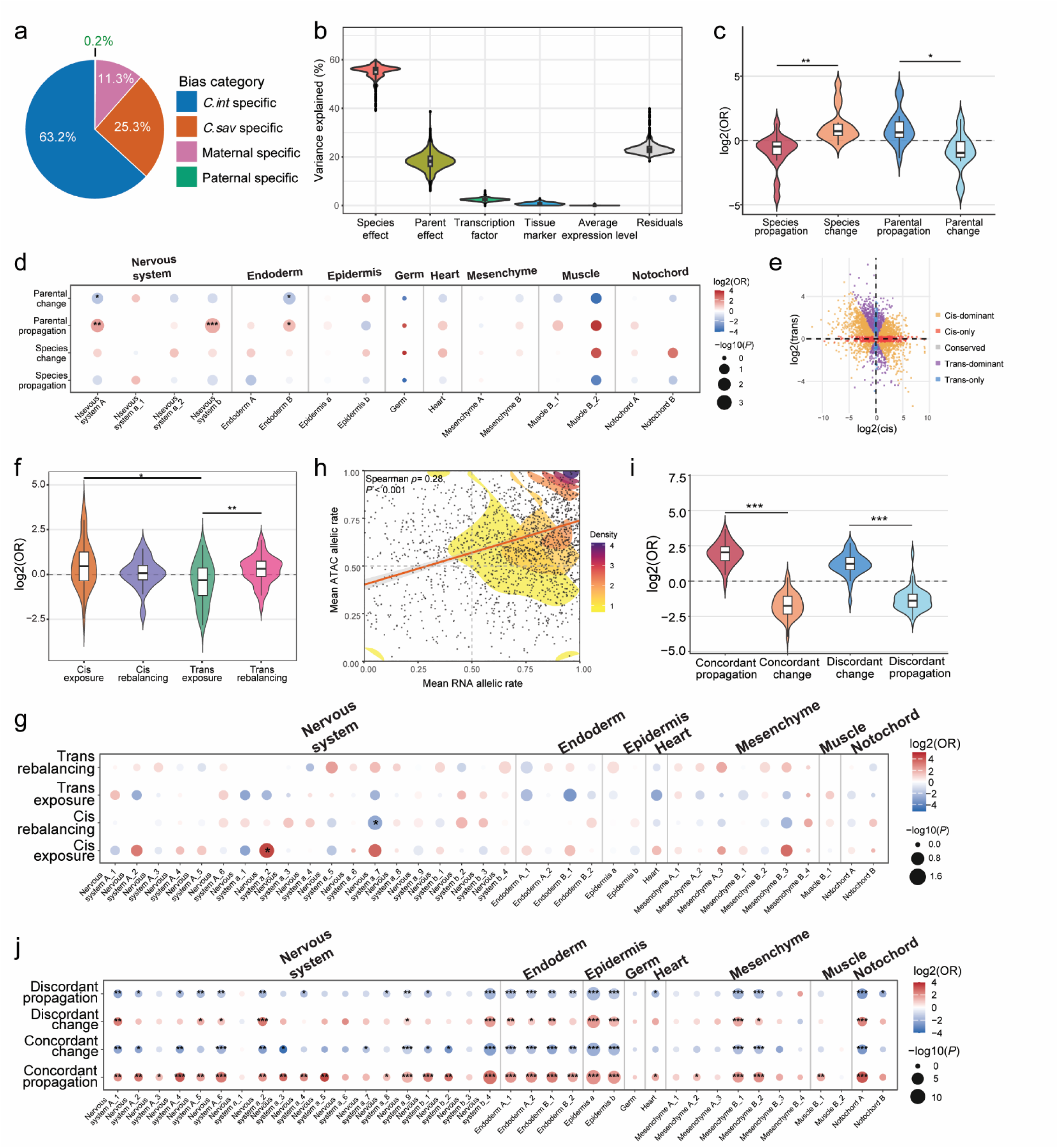
Regulatory architecture is associated with distinct allelic fates during lineage progression. **a,** Relative contributions of species-specific and parent-of-origin-associated biases at the mid-gastrula stage, determined using reciprocal hybrid crosses (n = 470 genes). **b,** Variance-partitioning analysis of allelic-rate variation at mid-gastrula. Species effect, parent-of-origin effect, transcription-factor status, and tissue-marker status were included as binary random effects, whereas average expression level was included as a continuous fixed effect. **c,** Association of species-specific and parent-of-origin-associated effects with subsequent allelic fate. Violin plots show log2 odds ratios (ORs) for enrichment of each regulatory class among propagated versus changed allelic states. Change includes switched, attenuated, and exposed transitions. Species propagation *vs*. Species change: *P* = 0.0025. Parental propagation *vs*. Parental change: *P* = 0.016. **d,** Lineage-resolved enrichment of species-specific and parent-of-origin-associated effects among propagated and changed allelic states. Color denotes log2(OR), and dot size denotes -log10(*P*). **e,** Cis- and trans-regulatory decomposition at the mid-tailbud stage based on parental expression divergence and allelic imbalance in the hybrid (n = 3,000 genes). Genes are classified as cis-dominant, cis-only, conserved, trans-dominant, or trans-only according to the relative contributions of cis- and trans-regulatory effects. **f,** Association of cis- and trans-regulatory classes with allelic exposure and rebalancing at the mid-tailbud stage. Violin plots show log2(ORs) for cis-associated exposure, cis-associated rebalancing, trans-associated exposure, and trans-associated rebalancing. Exposure denotes balanced to maternal/paternal transitions; rebalancing includes attenuated and buffered transitions. Cis exposure *vs*. Trans exposure: *P* = 0.036. Trans rebalancing *vs*. Trans exposure: *P* = 0.0081. **g,** Lineage-resolved enrichment of cis- and trans-associated exposure and rebalancing. Cis-associated genes include cis-only and cis-dominant classes; trans-associated genes include trans-only and trans-dominant classes. Color denotes log2(OR), and dot size denotes -log10(*P*). **h,** Correlation between mean RNA and chromatin-accessibility allelic rates at the mid-tailbud stage (n = 3,192 genes). Contours indicate local gene density, and the orange line indicates the fitted trend. Spearman’s correlation coefficient (ρ) and *P* value are shown. **i,** Association between RNA-ATAC concordance and subsequent allelic fate. Violin plots show log2(ORs) for concordant or discordant allelic regulation among propagated and changed states. Concordant genes show RNA and ATAC allelic biases in the same direction, whereas discordant genes show opposing allelic biases. Change includes switched, attenuated, and exposed transitions. Concordant propagation *vs*. Concordant change: *P* = 9.46E-08. Discordant propagation *vs*. Discordant change: *P* = 1.10E-06. **j,** Lineage-resolved enrichment of concordant and discordant RNA-ATAC regulation among propagated and changed allelic states. Color denotes log2(OR), and dot size denotes -log10(*P*). For **d, g,** and **j**, enrichment *P* values were calculated using Fisher’s exact tests. For **c, f,** and **i**, group differences were assessed using two-sided Mann-Whitney *U* tests. * *P* < 0.05, ** *P* < 0.01, and *** *P* < 0.001.

These regulatory-source classes were next compared with subsequent allelic behavior across outgoing lineage transitions. Odds ratio analysis showed opposing developmental outcomes for the two regulatory sources: species specific biases were preferentially associated with transitions involving changes in allelic state, whereas parent of origin associated biases were preferentially associated with propagated transitions (Fig. 3c). Tissue resolved analysis further showed that these relationships were distributed across multiple developmental lineages (Fig. 3d and Extended Data Table 9). Thus, parent of origin associated biases showed greater allelic memory, whereas species specific biases were more frequently remodeled as lineages progressed, showing that developmental trajectories differentially interpret regulatory differences according to their origin. The greater turnover of species-specific effects is consistent with the modular and context-dependent deployment of cis-regulatory differences across changing lineage-specific trans environments^25^.

We next dissected cis- and trans- regulatory divergence at the mid-tailbud stage, where matched transcriptomes from both species and viable hybrids enabled direct comparison of interspecies and allelic expression differences (Methods, Fig. 3e). Among 3,000 genes, cis-only effects accounted for 17.2% and trans-only effects for 6.9%, whereas approximately 74.6% showed combined cis and trans contributions. Within the combined class, cis- and trans- effects either acted in the same direction to reinforce species differences or in opposite directions, consistent with compensatory regulatory divergence (Extended Data Fig. 11)^24,26^. The remaining 1.2% showed no significant regulatory difference. These results indicate that allelic imbalance arises from both allele-intrinsic sequence divergence and differences in regulatory environment, with cis effects representing the stronger global component at this stage. We grouped cis-only and cis-dominant genes as cis-associated, and trans-only and trans-dominant genes as trans-associated, then examined how these regulatory classes were distributed across lineage-transition fates (Fig. 3f,g and Extended Data Table 9). Cis-associated divergence was preferentially linked to exposed transitions, in which a balanced allelic state in the parent lineage became biased in the descendant lineage. This pattern is consistent with a model in which lineage-specific regulatory programs activate divergent parental enhancers unequally in the shared hybrid environment. By contrast, trans-associated divergence was more often linked to rebalancing transitions, including attenuation or buffering of allelic bias. This pattern is consistent with parental trans-regulatory differences being reduced in the shared cellular environment of the hybrid embryo^27^. Thus, cis- and trans- architecture contains information about its subsequent allelic behavior during lineage progression.

Because chromatin accessibility can precede transcription and mark primed or maintained regulatory states^28-31^, we next asked whether allele-specific chromatin support predicted allelic memory. We utilized mid-tailbud scATAC-seq data and compared allele-specific RNA bias with allele-specific accessibility at linked regulatory peaks (Methods, Fig. 3h). RNA-ATAC comparison showed that chromatin support for allelic RNA bias was direction-dependent (Extended Data Fig. 12). Maternal-biased RNA genes showed high concordance with accessibility across tissues, with 67.1-79.1% retaining the same allelic class. By contrast, paternal-biased RNA genes showed lower concordance, with only 38.7-50.5% matching the accessibility class and 49.5-61.3% showing discordance. Thus, allele-specific chromatin accessibility contributes to transcriptional allelic imbalance, but its relationship to RNA bias differs between maternal- and paternal-biased states. Linking RNA-ATAC concordance to matched lineage transitions showed that concordant allelic imbalance was enriched among propagated transitions, whereas discordant states were enriched among stage changing transitions (Fig. 3i, j and Extended Data Table 9). Thus, chromatin-supported allelic imbalance is more likely to be remembered across lineage progression, whereas discordant RNA and chromatin bias marks a more labile regulatory state. This result adds a mechanistic layer to our framework, indicating that allelic bias is more likely to persist when transcriptional asymmetry is accompanied by concordant chromatin accessibility.

Together, these analyses support a model in which embryonic lineage progression acts as a regulatory sorting process. In this model, the molecular origin and chromatin support of allelic divergence help determine whether it is retained as developmental memory, rebalanced within the shared hybrid environment or remodeled as lineages diversify. Thus, developmental context may convert divergent regulatory inputs into distinct allelic outcomes, a process likely shaped by the lineage-specific deployment of embryonic GRNs^24,32,33^.

### GRN hierarchy links regulatory architecture to allelic fate

The preceding analyses suggested that developmental context helps sort allelic states during lineage progression. We next examined whether this sorting is organized by GRN hierarchy. Because developmental GRNs link upstream lineage-specification factors to downstream differentiation and target genes, their hierarchical organization provides a natural framework for assessing where allelic divergence becomes visible. Using published lineage-specific regulatory cascades^18^, we assigned genes to upstream, intermediate, downstream or target positions and quantified the fraction of expressed lineage-stage contexts in which each gene showed allele-specific activity (Methods). This allelic-divergence score provided a network-level measure of how often each regulatory node displayed allelic imbalance during development. Across endoderm, mesenchyme and peripheral nervous system lineages, allelic divergence generally increased along the GRN hierarchy, from upstream regulators to downstream regulators and target genes (Fig. 4a and Extended Data Fig. 13). Upstream regulators consistently showed the lowest divergence fractions, whereas target genes showed the highest. This pattern was most pronounced in endoderm but was also significant in mesenchyme and PNS, indicating that the hierarchical distribution of allelic divergence extends across multiple developmental lineages. Thus, GRN hierarchy is associated with where allelic divergence becomes developmentally manifest, with core upstream regulators showing lower divergence than downstream regulatory and target-gene layers.

**Figure 4.**
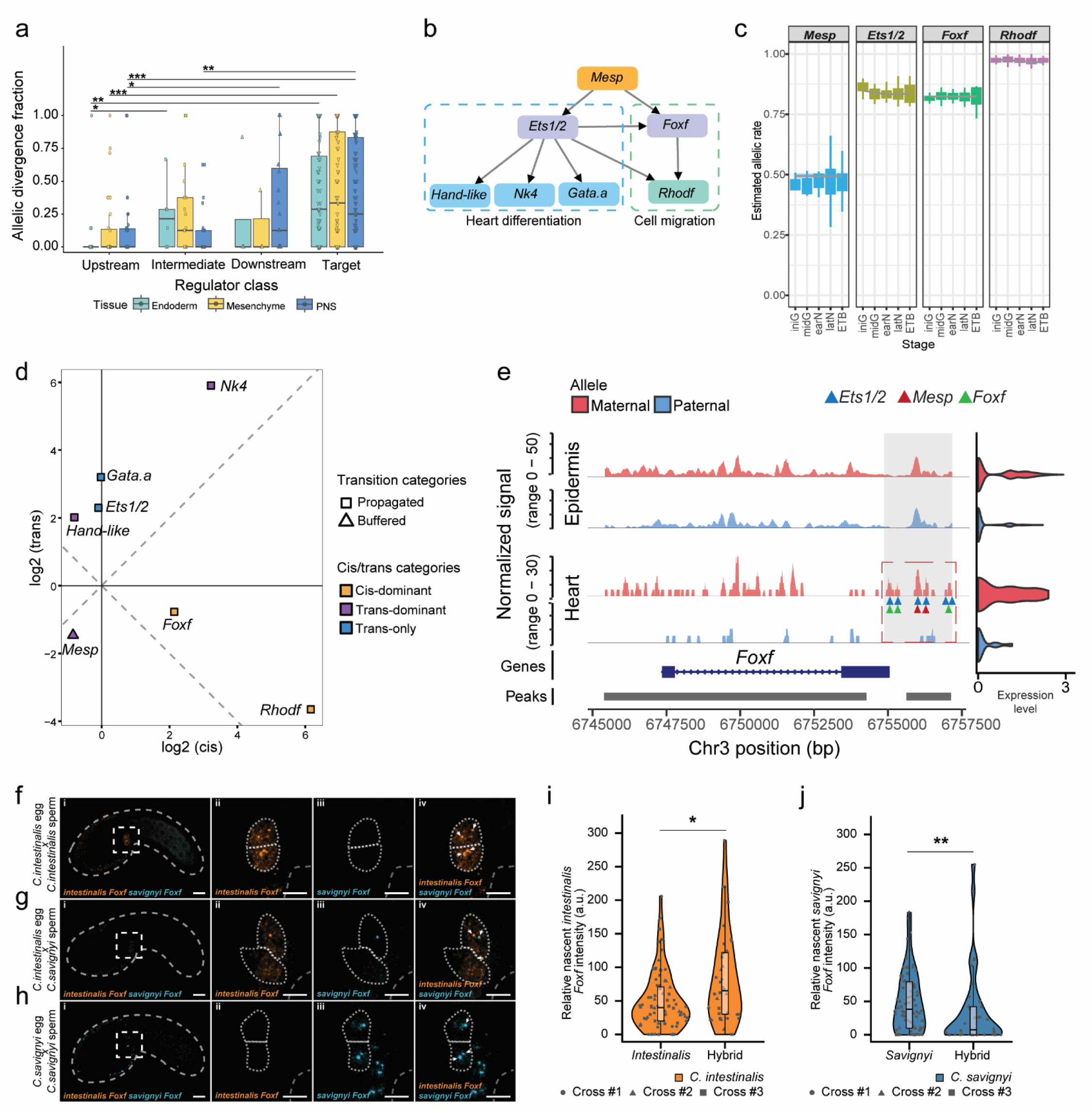
Cardiopharyngeal GRN architecture links network hierarchy to regulatory divergence and allelic behavior. **a,** Allelic divergence varies with hierarchical position in developmental gene networks across representative tissues (Endoderm, n = 119 genes; Mesenchyme, n = 134 genes; PNS, n = 236 genes). Boxplots are colored by tissue. PNS: peripheral nervous system. Endoderm: upstream *vs.* intermediate: *P* =0.034; upstream *vs.* target: *P* = 0.0013. Mesenchyme: upstream *vs.* target: *P* = 5.43E-04. PNS: upstream *vs.* downstream: *P* = 0.034; upstream *vs.* target: *P* = 3.96E-05; intermediate *vs.* target: *P* = 0.001. **b,** Schematic of the compact *Ciona* cardiopharyngeal GRN. The lineage initiator *Mesp* activates downstream regulators including *Ets1/2*, which in turn regulates the differentiation branch (*Hand-like*, *Nk4*, *Gata.a*) and the migration branch (*Foxf* and *Rhodf*). Arrows indicate activating regulatory relationships. **c,** Estimated allelic-rate dynamics of representative cardiopharyngeal GRN genes across developmental stages. **d,** Cis- and trans- regulatory decomposition of representative GRN genes at the mid-tailbud stage. The x axis indicates cis effects, and the y axis indicates trans effects. Colors denote regulatory classes, and symbol shape indicates lineage-transition fate category. **e,** Allele-resolved chromatin accessibility at the *Foxf* locus in epidermis and TVCs at the mid-tailbud stage. *C. intestinalis* (maternal) and *C. savignyi* (paternal) allele specific accessibility are shown separately for each cell type, with the highlighted region indicating putative regulatory regions near *Foxf* transcription start site. Marginal density plots at right summarize *Foxf* expression levels in epidermis and TVCs. Gene structure and accessible peaks are shown below. Triangles indicate predicted binding motifs for *Ets1/2*, *Mesp*, and *Foxf* within accessible regions. **f-h,** Allele-specific HCR detection of nascent *FoxF* transcription at the mid-tailbud stage in embryos from *C. intestinalis* egg × *C. intestinalis* sperm (**f**), *C. intestinalis* egg × *C. savignyi* sperm (**g**), and *C. savignyi* egg × *C. savignyi* sperm (**h**) crosses. In each row, (i) shows an embryo overview, with the dashed box indicating the region enlarged in (ii–iv); (ii) shows the *C. intestinalis Foxf* channel (orange); (iii) shows the *C. savignyi Foxf* channel (cyan); and (iv) shows the merged allele-specific signals. White arrows indicate nascent transcription foci. Dotted outlines delineate trunk ventral cells, and dashed outlines indicate embryo boundaries. Scale bars, 10 μm. **i,** Quantification of nascent *Foxf* transcription from the *C. intestinalis* allele in pure *C. intestinalis* and hybrid embryos. Statistical significance was assessed using a two-sided Mann-Whitney *U* test (*C. intestinalis*, n = 72 alleles; hybrid *C. intestinalis* allele, n = 36 alleles; *P* = 0.011). **j,** Quantification of nascent *Foxf* transcription from the *C. savignyi* allele in pure *C. savignyi* and hybrid embryos. Statistical significance was assessed using a two-sided Mann-Whitney *U* test (*C. savignyi*, n = 76 alleles; hybrid *C. savignyi* allele, n = 36 alleles; *P* = 0.0084). Signal intensity for each allele was normalized to the average total nascent *Foxf* fluorescence intensity per trunk ventral cell in same-species embryos in **i,j**. Points represent individual alleles, with shapes indicating independent crosses. \**P* < 0.05; \*\**P* < 0.01; \*\*\**P* < 0.001.

To examine this principle within a defined developmental network, we focused on the cardiopharyngeal lineage. This lineage is controlled by a compact and well-characterized GRN^34-37^, allowing allelic divergence to be compared across defined upstream and downstream regulatory nodes. In *Ciona*, the lineage initiator *Mesp* establishes the cardiopharyngeal program, followed by activation of downstream regulatory cascades involving *Ets1/2*, *Foxf* and *Rhodf* (Fig. 4b). Hybrid embryos preserved this conserved program, but allelic divergence was not evenly distributed across this network. *Mesp* showed balanced allelic contribution in the lineage, consistent with low divergence at the upstream lineage-specification layer, whereas downstream regulators showed stronger and more context-dependent allelic imbalance. In particular, *Foxf* and *Rhodf* displayed pronounced allelic bias in downstream lineage states (Fig. 4c). Thus, allelic divergence within the cardiopharyngeal GRN followed a hierarchical pattern, with the upstream lineage-specification node remaining balanced while selected downstream nodes became increasingly divergent. Overlaying cis- and trans- regulatory architecture and lineage-transition fate onto the GRN further revealed distinct regulatory modes across network branches (Fig. 4d; Methods). *Mesp* showed a buffered transition state and trans-dominant regulatory classification, whereas downstream regulators largely propagated their allelic states through different regulatory architectures. The cardiogenic branch, including *Gata.a*, *Nk4* and *Hand-like*, was predominantly trans-associated, whereas the migration associated *Ets1/2-Foxf-Rhodf* cassette showed strong cis-associated effects at *Foxf* and *Rhodf*. Thus, downstream branches can propagate allelic divergence through distinct cis-trans architectures despite comparatively low divergence at the upstream specification node.

We next examined the regulatory basis of *Foxf* allelic bias across developmental contexts. *Foxf* is expressed across multiple embryonic tissues, including the migratory trunk ventral cells (TVCs), and showed consistent bias toward the *C. intestinalis* allele across cell types, although the magnitude of allelic imbalance varied among developmental contexts (Extended Data Fig. 14). Total *Foxf* expression in TVCs differed markedly between the two species, whereas hybrid expression remained intermediate but closer to the high expression *C. intestinalis* state and was dominated by the *C. intestinalis* allele (Extended Data Fig. 14c). Allele-resolved chromatin accessibility further supported a cis-regulatory contribution to *Foxf* allelic bias. Consistent with the expression data, TVCs associated accessible regions near *Foxf* showed allele-biased chromatin accessibility, whereas comparable imbalance was not observed in other tissues such as epidermis (Fig. 4e). The biased accessible region contained allele-informative sequence differences overlapping predicted binding motifs for cardiopharyngeal regulators, including *Mesp*, *Ets1/2* and *Foxf* itself, suggesting that local sequence divergence contributes to allele-specific activity of this putative regulatory element (Fig. 4e). Allele-specific HCR further validated biased *Foxf* transcription within the TVCs (Fig. 4f-4j). Moreover, transcription from the single *C. intestinalis Foxf* allele was elevated in the hybrid relative to an individual allele in pure *C. intestinalis* (Fig. 4i), suggesting partial compensation for the weak contribution of the *C. savignyi* allele.

Together, this GRN provides a mechanistic example of regulatory sorting within a defined developmental network. By integrating lineage-resolved allelic transitions, cis-trans classification and allele-specific chromatin accessibility, this analysis shows how regulatory divergence is differentially distributed across individual GRN nodes and functional branches. Core specification can remain comparatively conserved while quantitative regulatory divergence emerges at selected downstream nodes, providing a potential mechanism by which developmental programs accommodate molecular evolution without disrupting overall lineage identity. More broadly, our lineage-resolved analyses reveal that regulatory differences accumulated between species acquire distinct developmental trajectories, and that these trajectories are systematically associated with regulatory origin and GRN hierarchy (Fig. 5). The molecular origin of regulatory divergence therefore contains information about its subsequent developmental behavior as cells progress through lineage-specific regulatory environments. Thus, developmental trajectories and regulatory architecture jointly shape where and when regulatory divergence becomes manifest during embryogenesis.

**Figure 5.**
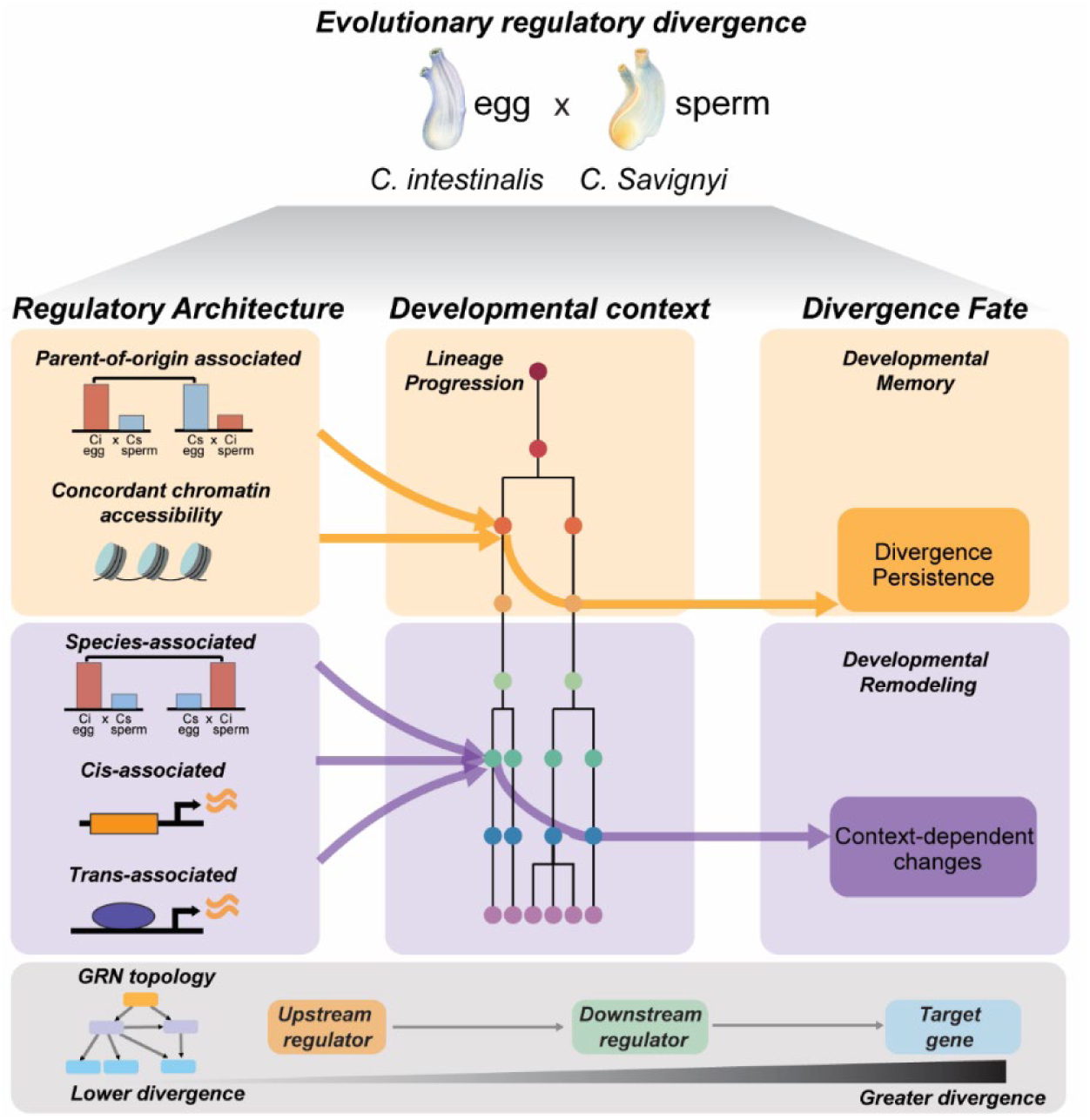
Developmental context and regulatory architecture shape the fate of evolutionary regulatory divergence. Schematic model summarizing how regulatory differences between *C. intestinalis* and *C. savignyi* are processed during hybrid embryogenesis. Parent-of-origin-associated biases and concordant allele-specific chromatin accessibility are associated with greater developmental persistence, whereas species-associated biases, cis-associated and trans-associated effects are more frequently associated with context-dependent changes in allelic state during lineage progression. GRN hierarchy provides an additional organizational axis, with lower allelic divergence at upstream regulators and greater divergence toward downstream regulators and target genes. Together, the model links molecular regulatory evolution to developmental outcome by showing how lineage context and GRN organization determine which evolved regulatory differences persist and which become developmentally manifest.

## Discussion

In summary, our study uses hybrid embryogenesis to examine how regulatory divergence is transmitted and accommodated across the lineage architecture of an animal embryo. By following species derived alleles at single cell resolution throughout *Ciona* development, we show that regulatory divergence is not transmitted as a fixed property of individual genes. Instead, allelic states are differentially retained or remodeled as cells progress through distinct developmental contexts. Their subsequent behavior is associated with developmental lineage, regulatory origin and GRN hierarchy. These findings provide a framework for understanding developmental system drift, in which conserved developmental outcomes are maintained despite divergence in their underlying regulatory mechanisms. This phenomenon is particularly relevant in ascidians, which retain highly similar embryonic development despite deep evolutionary divergence across lineages separated by approximately 390 million years^38^.

Developmental system drift has been proposed to arise through both compensatory evolution and the robustness of developmental regulatory networks^39-41^, with regulatory redundancy providing one mechanism that can increase such robustness. Our results suggest that several of these processes operate in the *Ciona* regulatory system. First, the asymmetric *Foxf* output suggests partial allelic compensation for the weak contribution of the *C. savignyi* allele. Allelic compensation has recently been demonstrated experimentally following allele specific enhancer perturbation^42^, providing a mechanistic precedent for our observation. Our results suggest that such compensation may also accommodate naturally evolved regulatory differences between species. Second, the cardiopharyngeal GRN provides evidence for branch specific network canalization. Across multiple tissues, allelic divergence increased toward downstream nodes, where additional cis regulatory elements may accumulate evolutionary changes^43,44^. In the cardiopharyngeal GRN, the migration associated branch showed strong cis associated divergence at *Foxf* and *Rhodf*, whereas the cardiogenic branch displayed a distinct regulatory architecture. This branch specific distribution suggests that regulatory divergence can be accommodated within individual GRN modules without propagating uniformly across the network, potentially reflecting network canalization. Regulatory redundancy may further contribute to this robustness by allowing individual regulatory elements to diverge while sufficient developmental output is maintained.

Together, our findings suggest how mechanisms implicated in developmental system drift may operate at different levels of regulatory organization. Allelic compensation can accommodate unequal output between divergent alleles, while network canalization and regulatory redundancy may limit the developmental consequences of regulatory divergence. Meanwhile, the accumulation of additional cis regulatory differences across successive GRN layers, together with their amplification through enhancer and chromatin responses, may help explain why divergence increases toward downstream GRN nodes. These complementary processes provide a mechanistic framework for how substantial regulatory evolution can accumulate while core developmental programs remain conserved. Over longer evolutionary timescales, such regulatory flexibility may provide substrate for more extensive GRN rewiring, redeployment or loss, as exemplified by the deconstruction of the ancestral tunicate cardiopharyngeal program in appendicularians^45^.

## Methods

### Ciona handling, collection, cross-species fertilization and dissociation of embryos

The adults of *C. intestinalis* and *C. Savignyi* were collected from the shore of California by M-Rep (San Diego, CA) and Marinus (Long Beach, CA) and shipped overnight to the laboratory. Gamete collection and egg dechorionation were performed as previously described^43^. Sperm was added to the eggs for 10 min. Then, the fertilized eggs were washed twice with filtered sea water. Except in the case of the larval stage, the same animal provided both the eggs and the sperm to lower the polymorphism rate for downstream analysis. Embryos were raised to different stages at 18°C according to a previously described method^44^. To prevent contamination, eggs and sperm were collected individually, and eggs were dechorionated. Dechorionated eggs were incubated for at least 1 h to confirm the absence of cleavage before insemination with sperm from the same or different species.

For each sample, 100 to 500 morphologically normal embryos were randomly picked and transferred into tubes pre-coated with 5% BSA in Ca^2+^-free artificial sea water with EGTA (Ca^2+^-free ASW, 10 mM KCl, 0.5M NaCl, 1.5 mM NaHCO_3_, 10 mM Tris-HCl pH8, 2.5mM EGTA). Embryos were immediately dissociated with 1% trypsin in Ca^2+^-free ASW, with the exception of the initial- and mid-gastrula stages, which were dissociated using 0.5% trypsin, and the larval stage, for which dissociation was performed using a mixture of 1% trypsin, 1 mg/mL collagenase, 0.5% pronase, and 0.5 mg/mL cellulase. Embryos were pipetted for 5 to 20 minutes depending on the stage to achieve the dissociation into individual cells. Then, the digestion was quenched by adding 20% fetal bovine serum (FBS) or 20% FBS with 2mg/ml glycine for larval stages. Cells were collected by centrifugation at 4 °C at 200g for 4 min and then resuspended in ice-cold Ca^2+^-free ASW containing 0.5% bovine serum albumin (BSA). The cells were then washed twice with 1% BSA in ASW and resuspended in 1% BSA in ASW without EGTA at a final concentration of 900-1500 cells/µl.

### Phalloidin staining and imaging

We fixed and stained embryos with alexa 488 phalloidin (ThermoFisher). Embryo are fixed in 100 mM HEPES [pH 7.0], 100 mM EGTA, 10 mM MgSO4, 2% paraformaldehyde (Electron Microscopy Sciences), and 300 mM dextrose, for 1 hour at room temperature, washed with PBT (0.1% Triton) 3 times, then incubated 24 hour in alexa 488 phalloidin. We collected confocal images on a Zeiss LSM 980 confocal microscope with a 10x/0.45 NA or 20×/0.6NA objectives.

### Single-cell RNA and ATAC sequencing library preparation

The cell concentration of each sample was checked by TC20 Automated Cell Counter to ensure it was within 1,000-2,000 cells per microlitre. For the scRNA-seq experiments, single-cell suspensions were loaded onto The Chromium Controller (10x Genomics). Cells were lysed, and cDNAs were barcoded and amplified with Chromium Single Cell 3′ Library and Gel Bead Kit v3 (10x Genomics) following the instructions of the manufacturer. For the single-cell multiomic experiments, isolated nuclei were processed using the Chromium Next GEM Single Cell Multiome ATAC + Gene Expression v1 assay with the Chromium Next GEM Chip J (10x Genomics), following the manufacturer’s instructions. For the single-cell ATAC-seq experiments, isolated nuclei were processed using the Chromium Single Cell ATAC Library and Gel Bead Kit (10x Genomics), following the manufacturer’s instructions. Illumina sequencing libraries were prepared from the scRNA-seq cDNA samples using the Nextera DNA library prep kit (Illumina). The scRNA-seq libraries were sequenced on Illumina HiSeq 2500 Rapid flowcells (Illumina) with paired-end 28 nucleotides (nt) + 125 nt reads following standard Illumina protocols. Raw sequencing reads were filtered by Illumina HiSeq Control Software and only pass-filtered reads were used for further analysis. Samples were run on both lanes of a HiSeq 2500 Rapid Run mode flow cell instrument. Base calling was performed by Illumina RTA version 1.18.64.0. BCL files were then converted to FASTQ format using bcl2fastq version 1.8.4 (Illumina). Reads that aligned to phix (using Bowtie version 1.1.1) were removed, as were reads that failed Illumina’s default chastity filter. We then combined the FASTQ files from each lane and separated the samples using the barcode sequences allowing one mismatch (using barcode_splitter version 0.18.2).

### Improvement of the *Ciona savignyi* genome assembly and ortholog gene mapping

An improved *Ciona savignyi* genome reference was generated using Dovetail Genomics Hi-C scaffolding^45,46^ and transcriptome-supported annotation files. Briefly, chromatin was crosslinked, digested, proximity-ligated and sequenced using the Dovetail Hi-C LinkPrep workflow. A draft *C. savignyi* genome assembly downloaded from Ensembl CSAV2.0^47^ was used as the reference assembly for Dovetail scaffolding. The resulting Hi-C read pairs were mapped to the reference assembly to order and orient contigs into long-range scaffolds. The resulting improved *C. savignyi* genome assembly was 177,054,150 bp in length, contained 202 scaffolds and had an N50 of 12,321,149 bp.

To improve transcript annotation, we incorporated additional *C. savignyi* RNA-seq datasets generated from multiple developmental stages, including initial gastrula, mid-gastrula, neurula, early tailbud, mid-tailbud, late tailbud and larval stages. RNA-seq reads were quality-trimmed using TrimGalore and aligned to the new *C. savignyi* genome assembly using HISAT2^48^. Transcript models were assembled using StringTie^49^ and merged with the existing *C. savignyi* ANISEED^50^ annotation using StringTie merge. After merging, annotation features lacking defined strand information or gene identifiers were removed to generate a cleaned StringTie-augmented GTF annotation.

For hybrid single-cell RNA-seq analysis, we generated a combined two-species Cell Ranger reference containing both *C. intestinalis* and *C. savignyi* genomes. The *C. intestinalis* component was built using the KY 2019 genome FASTA and corresponding GTF annotation downloaded from the Ghost database^51^. The *C. savignyi* component was built using the new genome assembly and the merged Dovetail-ANISEED-StringTie GTF annotation. The combined reference was generated with Cell Ranger v3.1.0 using cellranger mkref, with the two species specified as separate genome components.

To define orthologous relationships between C. intestinalis and C. savignyi, we compared the protein-coding gene sets from the *C. intestinalis* KY 2019 annotation and our improved *C. savignyi* annotation using OrthoFinder v2.3.12^52^. This analysis generated a revised ortholog gene list between the two species, including one-to-one, many-to-one and one-to-many orthologous relationships. The full ortholog table is provided in Extended Data Table 4.

### Single cell RNA-Seq data clustering and visualization

For scRNA-Seq clustering and visualization, single-cell gene-expression matrices were analyzed using Seurat v4.3.0^53^. To enable joint analysis of hybrid cells together with published *Ciona intestinalis* single-cell datasets^18^, species-specific read counts were collapsed to orthologue-level gene counts. Specifically, for each matched C. intestinalis-C. savignyi orthologue, reads assigned to the two parental species were summed to generate a single total expression count for that orthologue gene. This merged orthologue-level count matrix was used as the input matrix for normalization, dimensionality reduction, clustering and visualization. Datasets were then integrated using the Seurat integration workflow to identify shared correlation structure across samples and developmental stages. The integrated expression matrix was scaled and used for principal component analysis, nearest-neighbour graph construction, clustering and low-dimensional visualization using UMAP. Hybrid cells were annotated by comparison with the published *C. intestinalis* reference annotations and by inspection of established lineage and cell-type marker genes^54^. Stage- and lineage-specific annotations were further refined manually when necessary to ensure consistency with known *Ciona* embryonic lineages. Allele-specific count matrices were retained separately and were used only for allele-resolved expression and allelic imbalance analyses.

### Single-cell multi-omics and ATAC-seq data processing, integration, clustering, and visualization

Single-cell multi-omics data were processed using Cell Ranger ARC v2.0.1 for samples collected at the late tailbud II, 18- h and 24-h larval stages. Reads were aligned to a custom hybrid *Ciona intestinalis-Ciona savignyi* reference generated from the *C. intestinalis* KY reference and the improved *C. savignyi* Dovetail assembly. The sample-specific Cell Ranger ARC outputs containing matched gene-expression and chromatin-accessibility profiles were imported into Seurat^53^ for downstream dimensionality reduction, clustering, integration, and visualization. Because gene-expression and chromatin-accessibility measurements were obtained from the same cellular barcodes, cell-type annotations derived from the gene-expression profiles were directly associated with the corresponding chromatin-accessibility profiles.

Single-cell ATAC sequencing data were processed using Cell Ranger ATAC v1.2.0 for samples collected at the mid tailbud stage. Reads were aligned to the same custom hybrid reference. Filtered peak-barcode matrices generated by Cell Ranger ATAC were analyzed using Signac^55^ and Seurat^53^. Chromatin accessibility quality-control metrics were evaluated using fragment size distribution and nucleosome signal. Accessible features were selected and multiple feature-selection thresholds (q0, q25, q50 and q75) were tested to evaluate the effect of feature selection on dimensionality reduction and clustering. For the q0 feature set, LSI components 2-30 were used for UMAP visualization, neighbor graph construction and clustering. The q0 Signac object was used for integration with the matched mid tailbud stage scRNA-seq dataset. To assign cell identities to scATAC-seq cells, the scATAC-seq dataset was integrated with the matched-stage scRNA-seq reference using Seurat label transfer. Variable features from the scRNA-seq reference were used for anchor identification, and canonical correlation analysis was used for cross-modality alignment. Prediction scores were inspected, and scATAC-seq cells with a maximum prediction score greater than 0.5 were retained for downstream visualization and comparison with the matched scRNA-seq annotations. For motif analysis, peak coordinates were converted to genomic ranges, and peak sequence information was obtained from the corresponding *Ciona* genome FASTA file. Position frequency matrices from the JASPAR2018^56^ CORE collection were retrieved using TFBSTools^57^ and used to construct a peak-by-motif matrix with Signac. Differentially accessible peaks between selected scATAC-seq clusters were identified using Seurat with logistic regression, including total peak counts as a latent variable. Motif enrichment was then performed on the top differentially accessible peaks.

### Data quality control and allelic rates estimation

For allele-resolved scRNA-seq analysis, cells with fewer than 200 detected genes were removed from downstream analysis, and genes detected in fewer than three cells were excluded. Raw FASTQ files were processed using the count pipeline in 10x Genomics Cell Ranger v3.1.0 with default settings to generate gene-barcode matrices for each sample. Reads were aligned to a species-aware combined reference generated from the *Ciona intestinalis* reference sequence obtained from the Ghost database^51^ and our improved *Ciona savignyi* genome assembly. Only reads that uniquely aligned to one species genome were retained for allele assignment. Allele-informative reads were assigned to either the *C. intestinalis* or *C. savignyi* allele.

For each gene within each developmental stage and cell-type context, we quantified the relative contribution of the two parental alleles using scDALI^20^. Genes with nonzero allele-informative coverage in more than 30 cells were retained. Cell-state covariates were defined from the first 20 principal components of the single-cell transcriptomic embedding. Allelic imbalance was then tested using the scDALI-Joint or Heterogenous model in scDALI, with the expected base allelic rate set to 0.5. Raw P values were adjusted for multiple testing using the Benjamini-Hochberg false discovery rate procedure. Genes with FDR-adjusted q values less than 0.01 were considered to show significant cell-state-associated allelic imbalance. For genes passing the q < 0.01 threshold, posterior allelic rates were estimated using scDALI interpolation with the same 20-principal-component cell-state representation. Posterior mean allelic rates and posterior variances were exported for downstream visualization and quantitative analysis of allele-specific expression dynamics across developmental cell states. For allele-resolved scATAC-seq analysis, a chromatin assay was constructed in Signac^55^ using the peak count matrix and fragment file generated from output file of Cell Ranger ATAC v1.2.0. Peaks detected in fewer than 10 cells were excluded, and cells with fewer than 200 detected peaks were removed. The nucleosome signal and transcription start site (TSS) enrichment score were calculated for each cell using the NucleosomeSignal and TSSEnrichment functions, respectively. The percentage of reads in peaks was calculated as the number of fragments overlapping called peak regions divided by the total number of fragments passing Cell Ranger ATAC filtering. Cells were retained for downstream analyses if they had between 1,000 and 30,000 total peak counts, more than 30% of reads in peaks, a nucleosome signal below 4, and a TSS enrichment score greater than 1.

We also used scDALI to quantify the relative chromatin accessibility associated with the two parental alleles. Allele-informative peaks located within the gene body or its 2-kb upstream region were retained and assigned to the corresponding orthologous gene based on ortholog gene mapping. These assigned peaks were then used to construct allele-specific peak count matrices for subsequent scDALI analysis. To obtain a common cell-state representation and facilitate comparisons between RNA- and ATAC-derived allelic rates, scATAC-seq data were integrated with scRNA-seq data at mid-tailbud stage. The first 20 principal components derived from the integrated dataset were used to represent cell state in the scDALI analysis. Allelic imbalance was tested using the Joint, Heterogeneous, and Homogeneous models implemented in scDALI, with the expected baseline allelic rate set to 0.5. Raw *P* values from each model were adjusted for multiple testing using the Benjamini-Hochberg false discovery rate procedure. Genes identified as significant by at least one of the three models, corresponding to the union of genes detected by the Joint, Heterogeneous, or Homogeneous model, at an FDR adjusted *q* value threshold of 0.05 were considered to exhibit significant allelic imbalance. For genes passing this threshold, posterior allelic rates were estimated using scDALI interpolation based on the same 20-principal-component cell-state representation. Posterior mean allelic rates and posterior variances were exported for downstream visualization and quantitative comparisons of allele-specific chromatin accessibility across cell states.

To minimize the potential contribution of maternally deposited transcripts to the analysis of allelic imbalance, we used a previously published bulk RNA-seq dataset generated from stage 0 unfertilized eggs^58^. Genes with detectable expression in unfertilized eggs were classified as maternally loaded transcripts and excluded from subsequent analyses, including Figures 1d-1h, 2, 3, and Extended Data Fig. 4, 6-12, 15.

### Reproducibility

Reproducibility was evaluated based on cell-type composition and allelic imbalance using three independently generated scRNA-seq datasets from the late tailbud II stage. We first compared the relative abundance of annotated cell types across the three replicates to assess the consistency of cell-type composition. Allelic rates were then estimated independently for each replicate. Pairwise comparisons were performed to evaluate the concordance of gene-level allelic rates among replicates, both across all cells and within individual tissues. The similarity of allelic rates between each pair of replicates was quantified using Spearman’s rank correlation. To evaluate whether differences in sequencing depth affected the estimation of allelic bias, we additionally performed a read-down sampling analysis. Each replicate were randomly down sampled to 100 million reads from raw bam files. Allele-specific expression matrices were reconstructed from the down sampled data, and the first 20 principal components were recalculated to represent cell state. Allelic rates were subsequently re-estimated, and pairwise Spearman correlation analyses were repeated at both the all-cell and tissue levels to assess the robustness of allelic imbalance estimates after controlling for sequencing depth.

### Allelic bias score analysis

To quantify the direction and consistency of allelic bias across tissues, we first calculated the mean allelic rate of each gene within each major tissue by averaging its cell-level allelic rates across all cells assigned to that tissue. The resulting tissue-level allelic rates were converted into directional scores: values greater than 0.5 were assigned a score of +1, indicating maternal bias; values below 0.5 were assigned a score of -1, indicating paternal bias; and values equal to 0.5 retained a score of 0, indicating balanced allelic activity. For each gene, an allelic bias score was calculated as the mean of these directional scores across all available tissues. The resulting score ranged from -1 to +1. A score of +1 indicated consistent maternal bias across all examined tissues, whereas a score of -1 indicated consistent paternal bias. Intermediate scores indicated that the direction of allelic bias varied among tissues, with positive values reflecting a greater prevalence of maternal bias and negative values reflecting a greater prevalence of paternal bias. Genes with scores of +1 and -1 were classified as stable maternal and stable paternal bias, respectively, whereas genes with intermediate scores were classified as switching allelic bias.

### Allelic bias markers and Gene Ontology enrichment analysis

Tissue-associated allelic bias markers were identified from the gene-by-cell allelic rate matrix. Differential allelic bias analysis was performed separately for each cell type using an adjusted *P* value threshold of 0.05 and a minimum allelic-rate difference of 0.25. Candidate markers were further filtered according to their median allelic rate within the corresponding cell type. Genes with a median allelic rate greater than 0.5 were classified as maternal-bias markers, whereas genes with a median allelic rate below 0.5 were classified as paternal-bias markers. Gene Ontology (GO) enrichment analysis was performed separately for the maternal- and paternal-bias marker sets identified in each tissue. *Ciona* ortholog genes were mapped to their corresponding human orthologs, and GO Biological Process (BP) enrichment was conducted against the human annotation database using the clusterProfiler R package^59^. GO terms with Benjamini-Hochberg-adjusted *P* values below 0.05 were considered significantly enriched. For visualization and interpretation, significant GO terms were manually grouped into broader functional categories based on their descriptions, such as development, metabolism, and cell adhesion.

### Lineage tree mapping and reconstruction

Hybrid scRNA-seq data from different developmental stages and tissues were mapped to previously published lineage-resolved reference datasets^18^ using the anchor-based integration and label-transfer workflow implemented in Seurat. Cells with a maximum prediction score greater than 0.5 were assigned to the corresponding reference lineage node. Cells with prediction scores of 0.5 or lower were considered uncertain and were assigned based on their positions in the integrated clustering, according to the annotated lineage node with which they co-clustered. Parent-child relationships among the mapped hybrid lineage nodes were defined according to the developmental lineage edges reported in the published reference lineage tree^18^. For analyses within each lineage node, only genes with a summed raw count greater than 30 across all cells assigned to that node were retained. To improve the robustness of downstream analyses, lineage nodes containing five or fewer cells were excluded. Based on the estimated allelic rates, genes with rates greater than 0.55 were classified as maternally biased, those with rates below 0.45 as paternally biased, and those with rates from 0.45 to 0.55, inclusive, as balanced. Changes in the allelic-bias state of the same gene across each parent-child lineage edge were then classified into five transition categories: switched, propagated, attenuated, buffered, and exposed. A transition from maternal to paternal bias, or vice versa, was classified as switched; maintenance of the same bias direction (maternal to maternal or paternal to paternal) was classified as propagated; a transition from a biased state to a balanced state was classified as attenuated; maintenance of a balanced state was classified as buffered; and a transition from a balanced state to either maternal or paternal bias was classified as exposed.

### Bias-persistence analysis

To assess differences in allelic memory across tissues and developmental lineages, gene-level changes in allelic bias between parent and daughter nodes were classified into five transition categories: switched, propagated, attenuated, buffered, and exposed. For each lineage node, allelic-bias persistence was quantified as the proportion of propagated transitions among genes classified as propagated, switched, or attenuated:

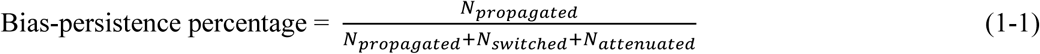

Higher percentage indicates that a greater proportion of genes retained the direction of allelic bias inherited from the parental node, whereas lower percentage indicates more frequent switching or attenuation of the pre-existing bias. Buffered and exposed transitions were excluded from this calculation because they did not represent the maintenance or alteration of a pre-existing directional allelic bias.

### Definition of species- and parent-of-origin associated biases

Species- and parent-of-origin associated allelic biases were classified using allele-specific rate matrices from the reciprocal crosses at mid-gastrula stage. An allelic rate greater than 0.5 was interpreted as maternal bias, whereas a rate below 0.5 was interpreted as paternal bias. Genes showing maternal bias toward the *C. intestinalis* allele in all evaluated cells of the forward cross and maternal bias toward the *C. savignyi* allele in all evaluated cells of the reciprocal cross were classified as exhibiting a maternal-specific bias. Conversely, genes showing paternal bias toward the *C. savignyi* allele in all cells of the forward cross and paternal bias toward the *C. intestinalis* allele in all cells of the reciprocal cross were classified as exhibiting a paternal-specific bias. Species-specific effects were defined according to consistency in the favored species across reciprocal crosses, regardless of parental origin. Genes showing maternal bias toward the *C. intestinalis* allele in the forward cross and paternal bias toward the *C. intestinalis* allele in the reciprocal cross were classified as exhibiting a *C. intestinalis*-specific effect. Similarly, genes showing paternal bias toward the *C. savignyi* allele in the forward cross and maternal bias toward the *C. savignyi* allele in the reciprocal cross were classified as exhibiting a *C. savignyi*-specific effect. Genes that did not satisfy any of these criteria were designated as unclassified.

For lineage-resolved analyses, the same classification criteria were applied to the mean allelic rate of each gene within each lineage node at mid-gastrula stage. Genes whose node-level mean allelic rates satisfied one of the definitions above were retained for downstream analyses.

### Definition of cis- and trans-associated regulatory categories

To quantify cis- and trans-associated regulatory classes, raw gene counts were first summed across all cells within each dataset to generate pseudobulk expression profiles for the *C. intestinalis* and *C. savignyi* alleles in the hybrid (HybridCi and HybridCs) and for the two parental species (ParentalCi and ParentalCs) at mid-tailbud stage. Each pseudobulk profile was independently normalized to counts per million (CPM). Genes were retained only when the combined raw count exceeded 30 for both hybrid alleles and also exceeded 30 across the two parental datasets:

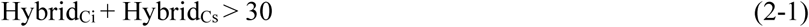

and

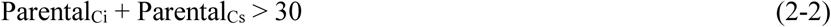

A pseudocount of 1 was added to each CPM value before ratio calculation. The cis effect was defined as the relative expression of the two alleles in the shared hybrid trans-regulatory environment:

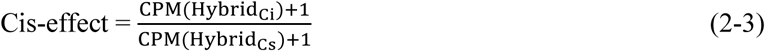

The total regulatory effect was defined as the expression ratio between the two parental species:

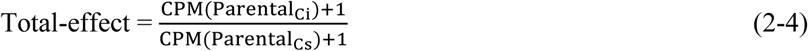

The trans-associated effect was calculated as the component of parental expression divergence not explained by the cis effect:

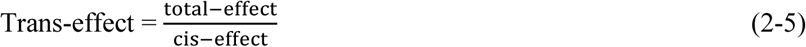

Ratios ranging from 0.8 to 1.2, inclusive, were considered unchanged. Genes were assigned to the following regulatory categories:

Conserved: both cis and total ratios were between 0.8 and 1.2.

Cis-only: the cis ratio was outside this interval, whereas the trans ratio remained between 0.8 and 1.2.

Trans-only: the cis ratio remained between 0.8 and 1.2, whereas the total ratio was outside this interval.

Cis-trans interaction: both cis and trans ratios were outside the 0.8-1.2 interval.

Genes classified as cis-trans interactions were further divided according to the relative magnitudes of the two effects. Genes with ∣log2(cis)∣>∣log2(trans)∣ were classified as cis-dominant, whereas genes with ∣log2(trans)∣>∣log2(cis)∣ were classified as trans-dominant. Genes with equal absolute cis and trans effects were designated as having equal contributions.

### Variance partitioning of allelic rate variation

Variance partitioning was performed using the variancePartition R package^60^. For each cell, estimated allelic rates across genes were logit transformed before model fitting. To accommodate allelic rates equal to 0 or 1, a small pseudovalue, 10^-3^, was added, and the transformed allelic rate was calculated as

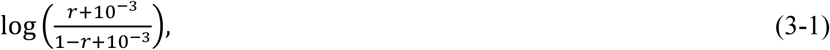

where *r* denotes the estimated allelic rate. Genes were matched to their corresponding annotation data, and categorical annotations were encoded as factors. A linear mixed-effects model was fitted independently for each lineage node using fitExtractVarPartModel. Average gene expression level was included as a fixed effect, whereas tissue-marker status, transcription-factor status, species-associated effect classification, and parent-of-origin-associated effect classification, RNA-ATAC concordance, and cis- and trans-regulatory classes were included as random effects. The fraction of within-cell allelic-rate variance attributable to each explanatory variable, together with the residual variance, was estimated for each cell.

### Classification of regulatory hierarchy and lineage specific target genes

Transcription factors were classified as upstream, intermediate, or downstream regulators based on their previously inferred activation timing in the lineage resolved regulatory cascades^18^. The transcription factor activation times were inferred from the temporal progression of gene expression along each developmental lineage and represented by the estimated expression onset parameter (time.on). For each lineage, the pseudotime range of the published regulatory cascade was divided into three equal intervals. Transcription factors with time.on values falling within the first, second, or third interval were classified as upstream, intermediate, or downstream regulators, respectively. Putative target genes were identified using the published lineage-resolved expression data^18^. Marker genes were calculated for each lineage at the larval stage and ranked according to their adjusted *P* values and log₂ fold changes. The top 50 marker genes for each lineage were selected, after which genes with annotated functional roles were retained as lineage-specific target genes.

### Quantification of lineage specific allelic divergence

For each gene within each lineage, raw maternal and paternal allele counts were summed across all cells. Allele-specific expression states were initially defined using the total gene count and the summed count for each allele. When the combined maternal and paternal count was below 10, both alleles were classified as off. For genes with a total count of at least 10, an allele was classified as on if its summed count was at least 5 and as off otherwise. To reduce the misclassification of lowly detected alleles as transcriptionally inactive, discordant allele states were further evaluated using the mean allelic rate estimated by scDALI. An initial maternal-on/paternal-off or maternal-off/paternal-on state was retained only when the mean allelic rate indicated near-monoallelic expression, defined as greater than 0.9 or less than 0.1, respectively. When the mean allelic rate was between 0.1 and 0.9, both alleles were reclassified as on. If a scDALI derived allelic rate was unavailable, the raw maternal allele fraction was evaluated using the same thresholds.

Each gene within lineage was consequently assigned to one of four states: on_on, in which both alleles were expressed; on_off, in which only the maternal allele was expressed; off_on, in which only the paternal allele was expressed; or off_off, in which neither allele was expressed.

For each gene within a tissue, the allelic divergence fraction across its constituent lineages was calculated as:

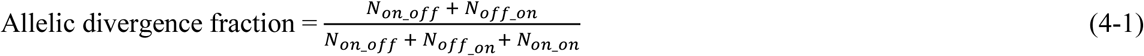

Lineage nodes in the off_off state were excluded because the gene was not detectably expressed. Values approaching 1 indicate a high prevalence of lineage-specific monoallelic expression and thus greater allelic divergence among lineages within the same tissue. Values approaching 0 indicate that both alleles are expressed across lineages, consistent with conserved biallelic expression.

### Classification of cardiopharyngeal regulatory modes and allelic transitions

For genes associated with the cardiopharyngeal gene regulatory network (GRN), cis- and trans- regulatory categories were assigned in TVCs at the mid tailbud stage using the criteria described above. To assess developmental changes in allelic bias, data from the late tailbud I and late tailbud II stages were combined into a single late tailbud group. Allelic transitions were then classified by comparing mean allelic rates in TVCs between the mid tailbud and late tailbud stages.

Allelic rates greater than 0.55 were classified as maternal biased, values below 0.45 as paternal biased, and values between 0.45 and 0.55 as balanced. Based on the change in allelic state between the two developmental stages, genes were classified as propagated when they retained the same directional bias, switched when the direction of bias changed between maternal and paternal, attenuated when an initially biased state became balanced, exposed when an initially balanced state became biased, or buffered when they remained balanced at both stages.

### Statistical analysis

Nonparametric comparisons between two independent groups were performed using the Mann-Whitney *U* test, whereas comparisons among three or more groups were performed using the Kruskal-Wallis test. Associations between variables were evaluated using Spearman’s rank correlation. Enrichment of intersections between gene categories was quantified using odds ratios, with statistical significance assessed using Fisher’s exact test. Where multiple hypotheses were tested, *P* values were adjusted using the Benjamini-Hochberg false discovery rate (FDR). *P* values, or adjusted *P* values where applicable, below 0.05 were considered statistically significant.

### In situ hybridization chain reaction (HCR) and image analysis

Embryos of *C. intestinalis, C. savignyi,* and the hybrid embryos from the *cross C. intestinalis eggs with C. savignyi sperm* were fixed at the mid-tailbud stage overnight at 4℃ using the fixative described by Treen and colleagues^62^, containing 100 mM HEPES, 500 mM NaCl, 1.75 mM MgSO4, 2 mM ethylene glycol bis (succinimidyl succinate), and 1% formaldehyde. The next day, the embryos were washed in PBST. They were progressively dehydrated with increasing concentrations of ethanol up to 80% ethanol and stored at -20 degrees Celsius. HCR was performed using the v3.0 RNAFISH HCR protocol following the recommendations of the manufacturer, Molecular Instruments, Inc. for sea urchin embryos except for the following steps. The embryos were blocked in the provided blocking buffer supplemented with 10µg/ml of salmon sperm DNA. After overnight hybridization with *C. intestinalis*-specific and *C. savignyi*-specific Foxf probes from Molecular Instruments (*intestinalis Foxf and savignyi Foxf*, respectively), the embryos were washed as recommended by the manufacturer. The hairpins conjugated to the fluorochrome Alexa Fluor 546 and Alexa Fluor 647 were denatured individually and combined with the amplification buffer. The amplification was performed for 1 hour and 30 minutes at 26℃. The embryos were then washed several times in 5x saline sodium citrate buffer (0.75M NaCl, 75mM Na_3_C_6_H_5_O_7_) with 0.1% Tween-20. They were then mounted on slides with RapiClear 1.49 (Sunjin Lab) and imaged with a Zeiss 880 confocal microscope.

The HCR signal intensity for *C. intestinalis Foxf* and *C. savignyi Foxf* in individual TVCs was measured by first manually segmenting the TVCs with FluoRender (v2.30). The two brightest 3D objects per TVC were then quantified as nascent transcription foci using custom scripts in ImageJ/Fiji (version 2.14.0/1.54j)^68^. Briefly, the background noise, defined as the mean pixel intensity of a trunk area without HCR signal, was first subtracted from the image. The image was converted to 8-bit. The signal of the *C. intestinalis* or *C. savignyi Foxf* transcript in individual TVC, based on the manually segmented mask, was then thresholded using the Renyi Entropy algorithm from the Auto Threshold plugin. 3D objects between 10 and 500 voxels were then segmented based on the Renyi Entropy threshold using the 3D Objects Counter plugin. If the Renyi Entropy threshold returned a value below 2, the gene was considered silent. The two brightest objects were assigned as the nascent transcription foci. The integrated density of the 3D object was used as the raw allele signal intensity before normalization. The images in Figure 4 are the sum-projection around the TVCs obtained using ImageJ/Fiji.

## Data availability

Single-cell RNA-Seq, multi-ome and ATAC-seq datasets, and assembled *C. savignyi* genome generated in this study are available in the NCBI Gene Expression Omnibus (GEO) under accession number GSE346238.

## Code availability

All code used in this study is available on GitHub at https://github.com/CaoLab-SysDevBio/Hybrid_project

## Acknowledgements

We thank the Princeton Lewis-Sigler Institute Genomics Core Facility, W. Wang, J. Wiggins and J. Miller for assistance with single-cell experiments; L. Parsons and J. Matese for contributions to the *C. savignyi* genome assembly; and the UT Dallas Office of Information Technology team for maintenance and support of the Ganymede 2 high-performance computing system. We thank all Cao group members for feedback.

## Contributions

J.L., L.L., M.L. and C.C. designed the study. J.L. and C.C. performed the computational analyses and contributed to data interpretation. L.L. performed the single-cell sequencing and HCR experiments. H.H. contributed to *Ciona* morphology imaging and manuscript revision. L.L. and L.R. contributed to HCR image analysis. J.L., M.L. and C.C. interpreted the results. J.L. and C.C. prepared the manuscript with input from all authors. C.C. supervised the project.

